# High throughput functional variant screens via in-vivo production of single-stranded DNA

**DOI:** 10.1101/2020.03.05.975441

**Authors:** Max G. Schubert, Daniel B. Goodman, Timothy M. Wannier, Divjot Kaur, Fahim Farzadfard, Timothy K. Lu, Seth L. Shipman, George M. Church

## Abstract

Tremendous genetic variation exists in nature, but our ability to create and characterize individual genetic variants remains far more limited in scale. Likewise, engineering proteins and phenotypes requires the introduction of synthetic variants, but design of variants outpaces experimental measurement of variant effect. Here, we optimize efficient and continuous generation of precise genomic edits in *Escherichia coli*, via in-vivo production of single-stranded DNA by the targeted reverse-transcription activity of retrons. Greater than 90% editing efficiency can be obtained using this method, enabling multiplexed applications. We introduce Retron Library Recombineering (RLR), a system for high-throughput screens of variants, wherein the association of introduced edits with their retron elements enables a targeted deep sequencing phenotypic output. We use RLR for pooled, quantitative phenotyping of synthesized variants, characterizing antibiotic resistance alleles. We also perform RLR using sheared genomic DNA of an evolved bacterium, experimentally querying millions of sequences for antibiotic resistance variants. In doing so, we demonstrate that RLR is uniquely suited to utilize non-designed sources of variation. Pooled experiments using ssDNA produced in vivo thus present new avenues for exploring variation, both designed and not, across the entire genome.

## Introduction

Constructing genotypes of interest and observing their effect on phenotype critically aids our understanding of genetics and genome function. As methods for editing genomes have progressed, this “reverse genetics” approach has expanded in breadth and scale, from knockout libraries^1^ to refactored genomes^2,3^. These experiments can now be performed within multiplexed pools, which allow an ever greater number of mutations to be explored across varied conditions. Critically, both creating genotypes and observing phenotype within pools has necessitated development of new techniques. Transposon insertions^4^, marked integrations^5^ and CRISPR-inhibition^6,7^ can create thousands of variants simultaneously within pooled experiments, and targeted sequencing of these elements enables pooled measurement of variant phenotypes. These advancements in experimental scale have fundamentally transformed our understanding of genome function^8^.

However, these current high-throughput genetic techniques remain limited, in that they typically introduce, ablate, or regulate kilobases of DNA to create variation. This contrasts with point mutations, which are ubiquitous in natural variation^9^, and are indispensable for engineering proteins^10^ and metabolic pathways^11^. While modifying kilobases of DNA can add and subtract functional elements such as genes and regulatory sequences from the genome, point mutations can alter the function of these elements, accessing a larger phenotypic landscape.

Point mutations and other precision edits can be performed by oligonucleotide recombineering, which creates precise genomic changes in many bacteria without incorporating selective markers or other large DNA sequences^12,13^ (Fig 1B). This technique enables multiple variants to be created simultaneously within pools^14^, but provides no means for determining the phenotypes of these mutations within pools. Therefore, genome-wide recombineering requires individual mutant clones to be isolated, genotyped, and phenotyped for causality to be established, severely limiting throughput.

**Figure 1:**
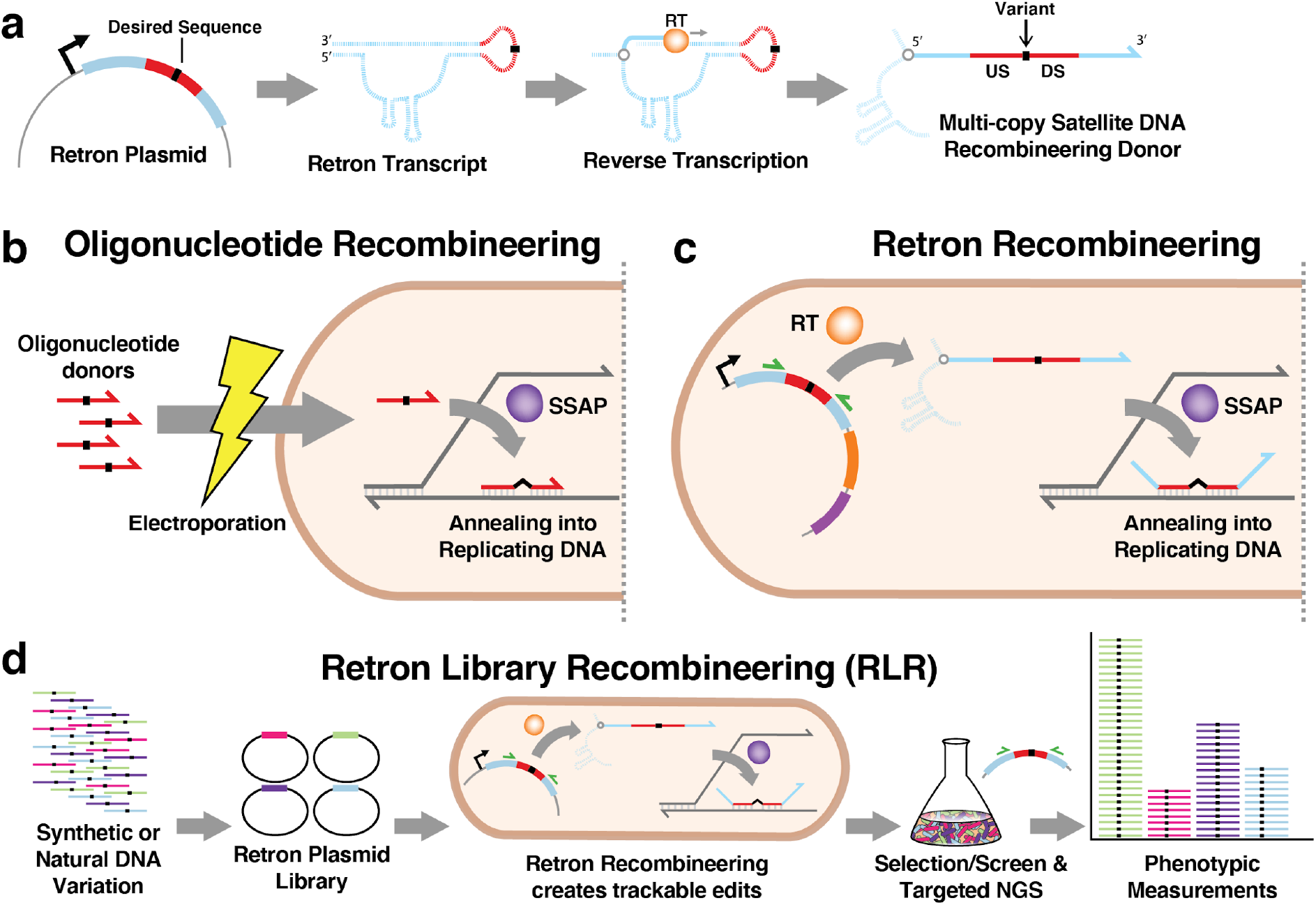
Graphical Introduction to Retron Library Recombineering. **1A** A retron contains a region msr/msd which undergoes targeted reverse-transcription, catalyzed by a retron reverse transciptase (RT), producing multi-copy satellite DNA. A novel sequence (red) introduced into retron becomes part of the reverse-transcribed product. When this sequence contains homology upstream (US) and downstream (DS) of a sequence alteration (black), multi-copy satellite DNA can be used as a single-stranded recombineering donor. RNA is depicted with dashed lines, whereas DNA is depicted with solid lines. **1B** Oligonucleotide recombineering is a technique in which synthetic oligonucleotide donors are introduced into bacteria and anneal to replicating DNA. This process is catalyzed by a Single-Stranded Annealing Protein (SSAP), and results in desired sequence alterations (black) being incorporated into the genome. **1C** Retron Recombineering is similar to oligonucleotide recombineering, but uses multi-copy satellite DNA produced by a retron as donor DNA, rather than transformed oligonucleotides. An SSAP is required for annealing of the donor, and retron Reverse Transcriptase (RT) is additionally required to create the donor. The retron plasmid remains in the cell, where it is available for targeted amplicon sequencing using complementary primers (green). **1D** Retron Library Recombineering (RLR) is a pooled functional genomics technique. Synthetic or natural DNA containing variation is used to construct a library of retron plasmids. Cells transformed with a given retron plasmid incorporate the variation within a retron donor, and the plasmids remain within cells as “barcodes” to distinguish mutant lineages. Next generation sequencing (NGS) of amplicons created using complementary primers (green) before and after a selection or screen reveals the abundance, and therefore the phenotypes, of individual variants.

We reasoned that it would be possible to transform recombineering into a method for pooled, genome-wide phenotypic measurement of large populations of precise genetic variants using bacterial retroelements known as retrons. Retrons are poorly-understood prokaryotic retroelements that undergo targeted reverse-transcription, producing single-stranded multi-copy Satellite DNA (msDNA)^15,16,17^. Previous work has shown that msDNA can function as a recombineering donor, creating specific edits in the genome^16,17^. This is accomplished by altering the retron sequence to produce single-stranded DNA containing a mutation of interest surrounded by homology to a targeted genomic locus (Fig 1A, 1C). Retron recombineering was used for genome modification in response to a stimulus, as a form of memory^17^, but editing rates far lower than canonical recombineering using electroporated DNA^14,18^ made it impractical for studying mutants of physiological interest.

Here we explored the factors which limit retron recombineering, and demonstrate significant improvements to its efficiency. We introduce Retron Library Recombineering (RLR), wherein a plasmid-encoded retron element creates specified mutations at high frequencies and remains in the cell to quantify mutants within multiplexed selections and screens via targeted sequencing (Fig 1D). Efficient editing and targeted amplicon sequencing of retron cassettes allows pooled phenotypic measurement of mutations across disparate genomic locations, including millions of retron cassettes blanketing the entire genome. We show that RLR is a generalizable, precision tool for multiplexed, high-throughput reverse genetics experiments.

## Results and Discussion

### Genotypic changes improve retron recombineering

Previous work has established that a recombineering donor can be introduced into the retron-derived msDNA, resulting in editing of the target locus when a single-stranded annealing protein (SSAP) such as Redβ of Enterobacteria phage λ is co-expressed^16,17^ (Fig 1C), but at a lower efficiency than when electroporated oligonucleotides are used as a donor. If retron recombineering could be made much more efficient, most cells bearing a given retron plasmid would contain the corresponding mutation. Thus, the retron cassette itself could serve as a “barcode” to identify mutants within mixed pools, enabling pooled screens of precisely-created mutant strains.

To measure editing, retron plasmids were constructed to confer drug resistance variants within the essential genes *rpoB* and *gyrA*, and co-expressed with Redβ (Methods). The fraction of cells edited after growth and induction of this system in batch culture for approximately 20 generations was measured by deep-sequencing the targeted locus, and by plating efficiency on the relevant antibiotic (Fig 2A, Fig S1C). Initially, less than ~0.1% of *E. coli* bearing the retron recombineering system incorporated the desired mutation (Fig 2B), replicating previous results^17^.

**Figure 2:**
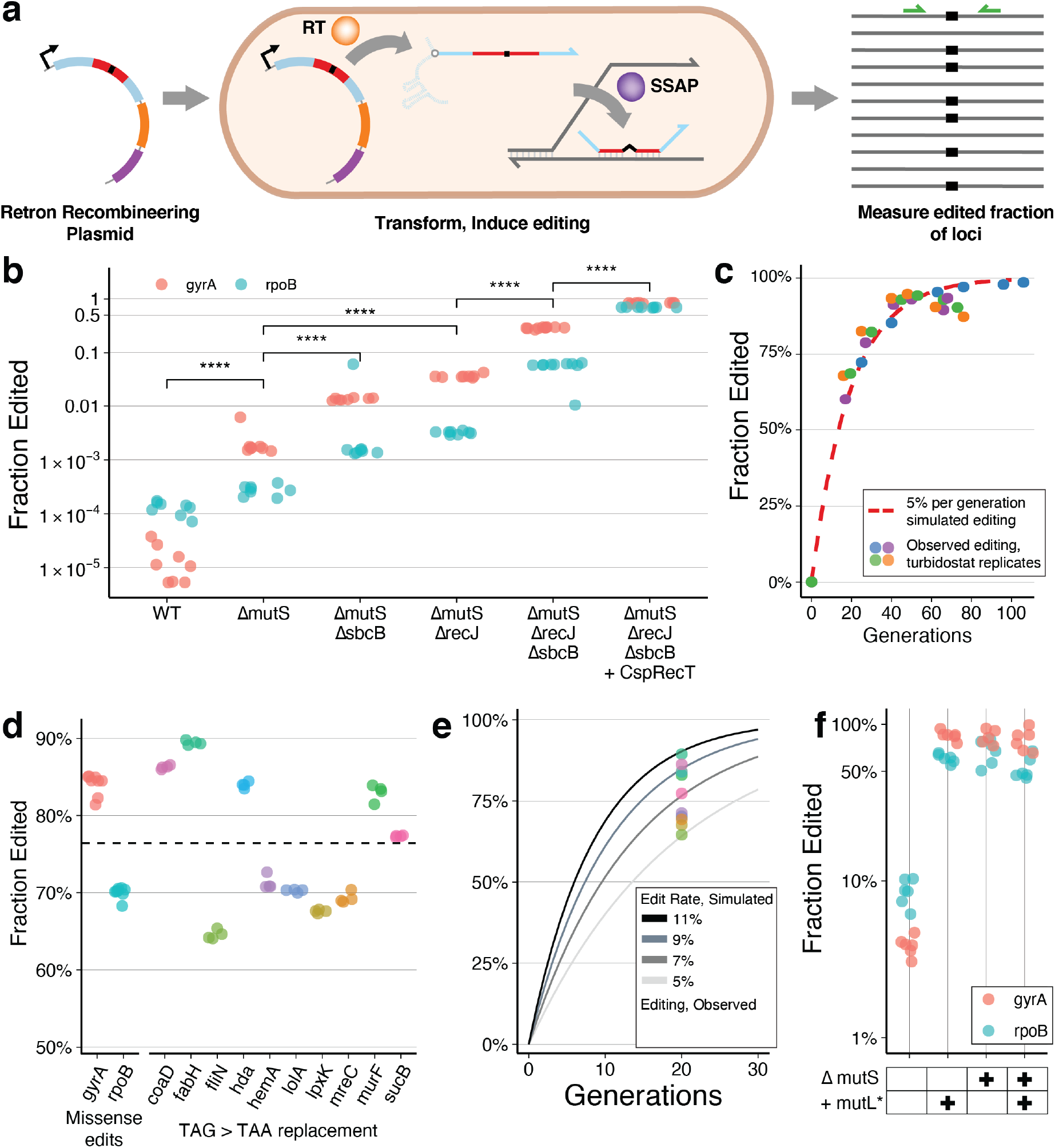
Editing the Genome using Retrons. **2A** Experiment measuring the edited fraction obtained for a desired mutation. A retron plasmid creates multi-copy single stranded DNA via a retron Reverse Transcriptase (RT). This DNA is incorporated into the genome via a co-expressed single stranded annealing protein (SSAP) as in Figure 1C. The edited fraction of resulting genomes is observed by amplicon Next Generation Sequencing (NGS), facilitated by primers targeting the locus (green). **2B** The effect of genotype on the edited fraction of loci, measured after induction of retron system and batch growth for approximately 20 generations. Colors differentiate experiments incorporating mutations at the *gyrA* and *rpoB* loci. Individual replicates are indicated with dots. “****” indicates p < 0.0001 resulting from a two-sided, unpaired parameteric t-test performed between genotype groups indicated with brackets. **2C** Δ*mutS* Δ*sbcB* Δ*recJ* cells were transformed with the *gyrA*-editing retron plasmid expressing Beta as an SSAP. Transformed cells were grown and induced continuously in 4 replicate turbidostats for 73 hours, sampling over time. The number of generations experienced by a given timepoint was inferred from the turbidostat growth record, and the edited fraction was determined by *gyrA* amplicon NGS for each sample. These data are shown alongside a simulated editing trajectory of an allele with neutral fitness effect, editing at 5% per generation. **2D** Edited fraction observed across alleles, after approximately 20 generations of induction and batch growth. A retron plasmid containing CspRecT as an SSAP was expressed in the Δ*mutS* Δ*recJ* Δ*sbcB* background in all cases. Results of individual replicates are shown as dots. Results for *gyrA* and *rpoB* missense alleles (Fig 2B) are shown again for comparison alongside TAG>TAA stop codon editing for 10 essential genes. The mean edited fraction achieved across the 12 loci, 76.4%, is indicated by dashed line. **2E** Mean edited fraction of the loci in Figure 2D are shown alongside a simulated editing trajectory of alleles with neutral fitness effect. This provides estimates of editing rates across alleles as within the range of 5-11% per generation. **2F** Retron plasmids with and without a dominant-negative MutL-E32K variant (*mutL*\*)^21^ were expressed in batch growth, in strains with and without inactivation of *mutS* (Δ*mutS*). Retron plasmids targeted the *gyrA* and *rpoB* loci and used CspRecT as an SSAP. Four replicate experiments per locus and condition are indicated with dots.

We first sought to improve this editing by investigating mismatch repair, which is known to interfere with oligonucleotide recombineering because the desired edit forms a genomic mismatch as a repair intermediate^12,19^ (Fig 1B, S1A). Inactivating mismatch repair by disruption of *mutS* improved the frequency of retron recombineering approximately 150-fold and 2-fold for *gyrA* and *rpoB* donors, respectively (Fig 2B). This difference in effect is explained by the known variation in sensitivity to mismatch repair among nucleotide mismatches^13^. Mismatch repair was therefore inactivated in subsequent experiments.

Additionally, previous studies have demonstrated that inactivation of exonucleases improves oligonucleotide recombineering by up to 3-fold^20,21^, and this effect has been shown for retron recombineering as well^22,23^. Inactivation of exonuclease genes *recJ* or *sbcB* (also known as *xonA*) individually provided benefit, and inactivation of both together increased the fraction of genomes edited during induction of retron recombineering in batch growth by 131 and 201-fold for the *gyrA* and *rpoB* donors, respectively (Fig 2B, S1C). We suspect that exonuclease inactivation results in a more dramatic benefit for retron recombineering, because recombineering donors are available continuously at low abundance, in contrast to canonical oligonucleotide recombineering, where donors are available transiently at high abundance.

Because editing occurs continuously in this system, we then explored monitoring the rate of editing over time, and whether prolonged editing produces a high fraction of edited cells. During continuous growth and induction of retron recombineering at the *gyrA* locus in a turbidostat, edited alleles accrued at a rate consistent with at least 5% editing per generation(Fig 2C, Fig S2). 73 hours of growth and induction across four replicate experiments resulted in edited fractions as high as 99%, with a mean of 92% (Fig 2C, Fig S1D).

### Optimization of RLR co-factors

We built a population genetics model simulating both editing and the abundance of edited populations to determine the effect of editing efficiency in multiplexed experiments (Fig S2A). We found that beneficial phenotypes like antibiotic resistance can be observed and quantified accurately with modest editing efficiency, because edited alleles will rapidly out-compete their non-edited counterparts once selection is applied (Fig S2C). Detrimental alleles, however, require more potent editing to observe even lethal effects (Fig S2D). Detrimental phenotypes also require persistent induction to be observed, whereas beneficial alleles can be observed after an initial pulse of induction (Fig S2B).

These considerations led us to seek further improvements in editing rate by exploring other SSAPs, the proteins required to catalyze recombineering by recruiting and annealing ssDNA to the replication fork^19^. While Redβ is the canonical SSAP used in *E. coli*, its distant relative CspRecT was found to improve recombineering efficiency in *E. coli*^24^. Replacing Redβ with CspRecT further increased the edited fraction observed in batch growth by nearly 3-fold to more than 12-fold for *gyrA* and *rpoB* donors, respectively(Fig 2B, S3B). Notably, requirements for inactivating mismatch repair and exonucleases are also somewhat relaxed when using CspRecT (Fig S3A).

To characterize improved editing across a sample of sequences and genomic regions, we altered TAG “amber” stop codons to TAA “ochre” variants, a target of intensive recombineering to produce highly altered genomes^25^. Altering 10 amber stop codons within essential genes by induction as before resulted in edited fractions from 65% to 89%(Fig 2D), consistent with editing rates in the range of 5-11% per generation(Fig 2E). The mean edited fraction across all loci examined was 76% (Fig 2D), exceeding the highest efficiency editing achieved using oligonucleotide recombineering, to our knowledge.

We also sought to remove the requirement for the mutagenic Δ*mutS* genotype for efficient editing. When a dominant-negative *mutL* allele^13^ (+*mutL*\*) is transiently expressed, the observed editing is equivalent to that observed in a Δ*mutS* background. This both minimizes the genotypic requirements in the parent strain and the duration of mutagenic mismatch-repair-deficient growth required (Fig 2F).

In summary, retron recombineering in the Δ*mutS* Δ*recJ* Δ*sbcB* background is thousands-fold improved over Wild Type, and can result in edited fractions greater than 90% with continuous editing. These improvements enable the presence of a retron cassette to identify its corresponding mutant strain within pooled experiments, and we proceeded to use this system for pooled experiments characterizing antibiotic resistance alleles. In tandem, we developed further improvements using the co-factors CspRecT and *MutL*\*. This system achieves higher editing rates and functions in a non-mutagenic strain background, enabling more demanding applications of RLR in the future.

### Generation of barcoded mutant pools and detection of phenotypes using RLR

With efficient retron recombineering established, targeted sequencing of retron cassettes can be used as a measure of a mutant’s abundance in a population, and therefore its phenotype within a pooled assay (Fig 1D). Antibiotic resistance is a growth phenotype of clinical importance, so we first sought to investigate antibiotic resistance mutations using RLR. We constructed a retron library conferring known *E. coli* rifampicin resistance mutations, known Mycobacterium tuberculosis rifampicin resistance mutations, and mutations affecting resistance to other antibiotics, and Neutral, deleterious, and lethal control mutations (Fig 3A). Redβ was expressed from the plasmid as an SSAP.

**Figure 3:**
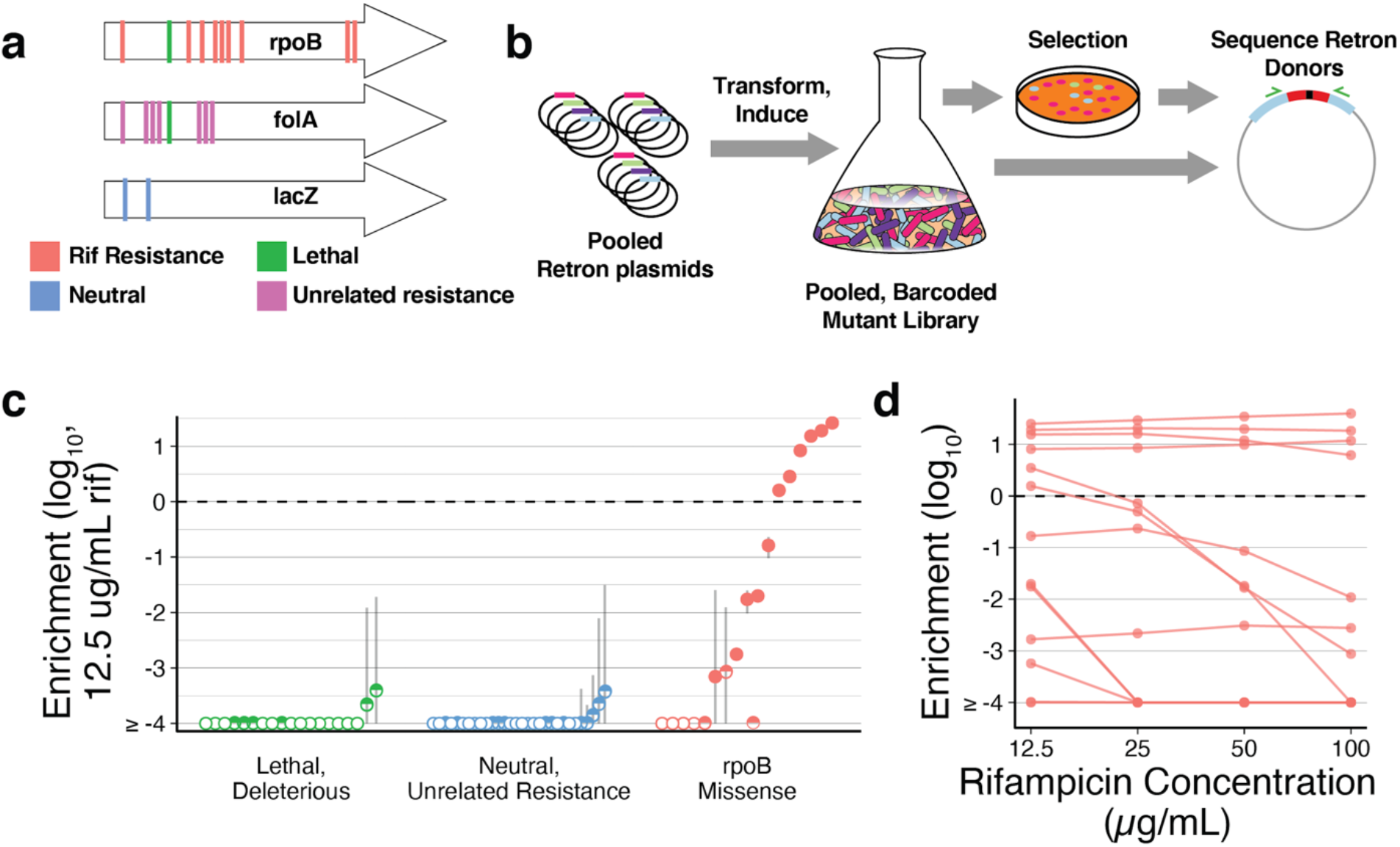
Pooled measurement of phenotypes using RLR. **3A** Example alleles contained in a synthetic pool. Both known and suspected rifampicin resistance alleles were specified in the pool, as well as resistance alleles for unrelated drugs, and control alleles expected to be neutral, lethal, or deleterious. For more details on edits specified, see Online Methods and Supplemental Table 2. **3B** Graphical representation of an RLR experiment to identify antibiotic resistance alleles. A pool of Retron plasmids conferring alleles of interest are transformed into cells. These transformants are induced, resulting in a pooled, barcoded mutant library. The library is subjected to selection, and the frequencies of retron donors are compared before and after treatment to calculate a resistance phenotype for each allele. **3C** RLR enrichment values of alleles, observed in a rifampicin treatment experiment. The median of three replicate experiments is indicated with a dot, and error bars are the Standard Error of the mean. A pseudocount of one is given to any allele not detected by sequencing after treatment, such that frequencies reported are a lower limit of detection in these cases. Unfilled points indicate when an allele was not detected among any of the three replicates after rifampicin treatment, and a half-filled point indicates when an allele was only detected in a subset of replicates. An enrichment value of zero is marked with a horizontal dashed line, indicating identical relative abundance before and after rifampicin treatment. **3D** For *rpoB* missense alleles, allelic enrichment across concentration of rifampicin is displayed. The median of three independent experiments is indicated with a dot, and lines connect an allele across concentrations.

Δ*mutS* Δ*recJ* Δ*sbcB E. coli* were transformed with this retron plasmid library, were induced in batch growth to acquire the desired mutations, and plated on solid medium with rifampicin. Sequencing retron donors from these samples before and after selection correctly identified known rifampicin resistance mutations^26^ by enrichment, while resistance alleles to unrelated drugs and other control alleles were depleted^7–9^(Fig 3C).

Mutations observed in rifampicin-resistant Mycobacterium tuberculosis^27^ did not confer detectable resistance in *E. coli*, with the exception of alleles altering the H526 residue of RpoB previously implicated in *E. coli* rifampicin resistance. This suggests some context-dependence of rifampicin resistance mutations across differing *rpoB* sequences.

A range of enrichment values was observed across resistance alleles, reflecting variation in the ability to grow under selection. Selection across rifampicin concentrations establishes inhibition curves for mutants within the pool (Fig 3D). In this way, RLR enables high throughput, pooled identification of antibiotic resistant alleles and facile characterization of their relative effects across a range of conditions.

### Quantitative pooled measurement of growth rate

Many phenotypes of interest produce small changes in fitness, rather than binary growth phenotypes^9,28^. Mutations producing small fitness improvements at low antibiotic concentrations can lead to the development of high-level antibiotic resistance^29^. To use RLR to quantify growth rate of mutants, we thus used growth in sub-inhibitory rifampicin as a model phenotype.

The pooled, barcoded mutant library constructed previously by RLR was grown at a sub-inhibitory concentration of rifampicin (5ug/mL), and samples were collected across multiple time points. Relative abundance of plasmids remained stable during transformation and induction of editing, but began to diverge once sub-inhibitory rifampicin was applied (Fig 4B). The degree to which resistant mutants outpace neutral controls is a quantitative measure of growth rate which correlates well to mutations when their growth is measured individually (Fig 4C, S4A). RLR, therefore, provides a quantitative growth rate measurement for alleles within a pool that is comparable to testing all mutants individually, but inherently more scalable.

**Figure 4:**
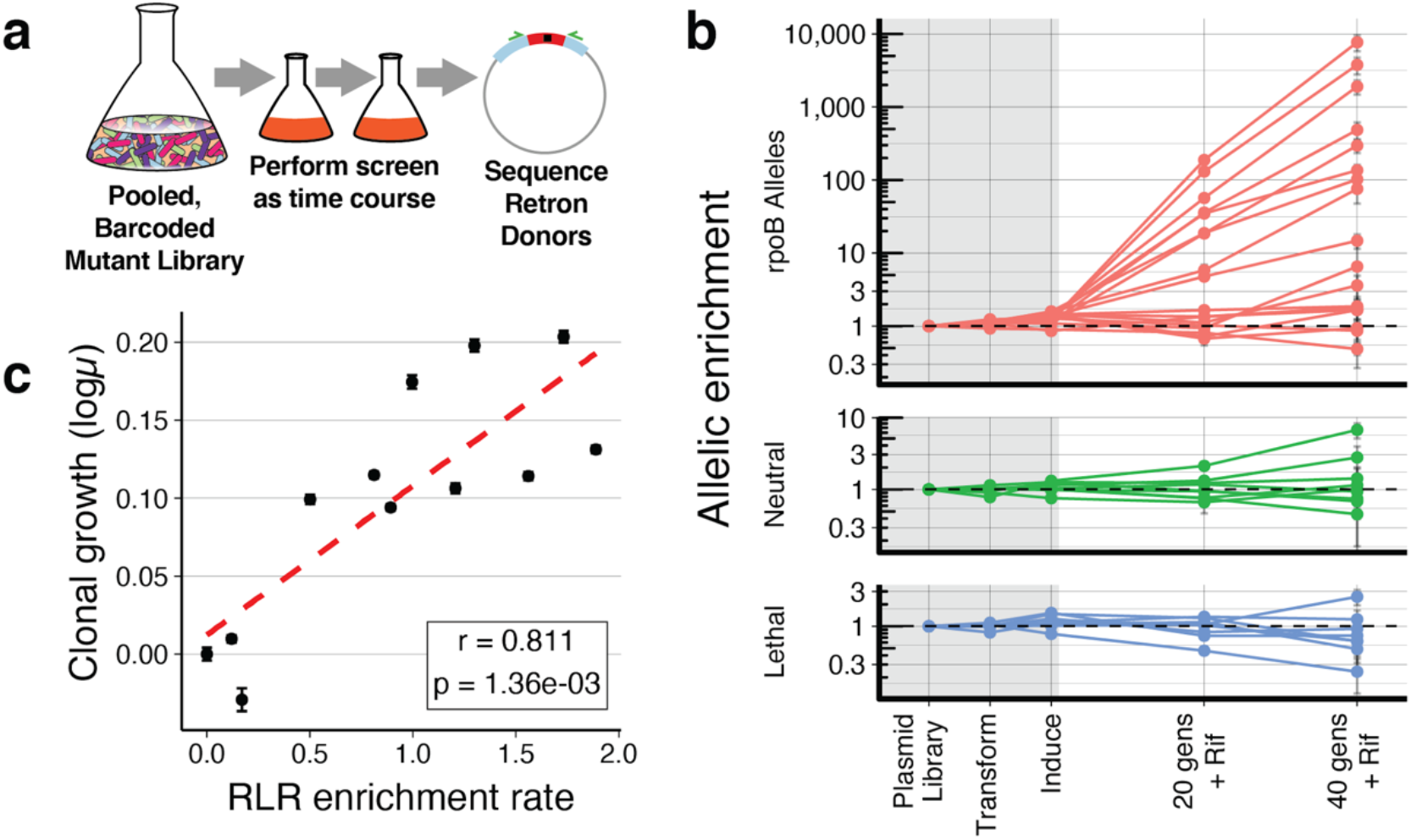
Quantitative measurement of phenotypes using RLR. **4A** Graphical representation of a quantitative RLR experiment. A pooled, barcoded mutant library constructed via RLR is subjected to a treatment time course, and the relative abundance of retron donors are measured over time. **4B** Frequencies of retron plasmids measured by targeted Next Generation Sequencing during an experiment are shown. Frequencies of each allele are normalized to the median of neutral controls within each timepoint, and to their starting abundance in the plasmid pool, for clarity. Frequency measurements within the grey rectangle occur during construction, transformation, and induction of the retron library. Measurements in the white area occur during growth in sub-inhibitory rifampicin, 5ug/mL. The mean across three replicates is indicated with a dot, and error bars are the Standard Error of this mean. Horizontal dashed lines indicate an allelic enrichment of 1, no change in frequency. **4C** A measure of growth rate for all alleles in the experiment can be determined by fitting an exponential curve to normalized frequency during the experiment, and reporting the the slope of normalized log10 allelic enrichment, over time. For a sample of 11 resistance mutants, the mean RLR slope across 3 replicate experiments is plotted on the X axis, and log10 of growth rates measured individually using classical methods are plotted on the y-axis. r is Pearson’s correlation coefficient between these two measures, and the p-value is the probability of these results given the null hypothesis of no correlation. Wild-type growth inferred from Neutral controls is plotted as an additional point. Error bars depict the standard error of clonal growth measurements, data visualized in Figure S4A

### RLR detects causal variants by using libraries prepared from evolved genomic DNA

Antibiotic resistance can evolve in the lab, producing strains with numerous mutations, able to grow in thousands-fold more concentrated antibiotic than their ancestors^30^. We reasoned that RLR could be used to determine the causal mutations leading to resistance in such isolates by using random fragments of their genomic DNA to construct an RLR library. Most fragments containing a variant should be capable of editing because we found little effect of mutation position within donor oligonucleotides (Fig S3C).

Genomic DNA from an *E. coli* strain highly resistant to the antibiotic Trimethoprim^30^ (TMP) was acoustically sheared into fragments, ligated to custom adapter sequences, and used to construct RLR libraries containing tens of millions of members in Retron plasmids expressing Redβ as an SSAP (Fig. 5A). Retron donors comprised sequences averaging approximately 100bp in length, providing over 50-fold coverage of the genome in unique fragments (Fig 5B, S6A, S6B), ensuring that variants present in the evolved genome are well-represented in the genomic RLR library. This library was then introduced into a Δ*mutS* Δ*recJ* Δ*sbcB* strain.

**Figure 5:**
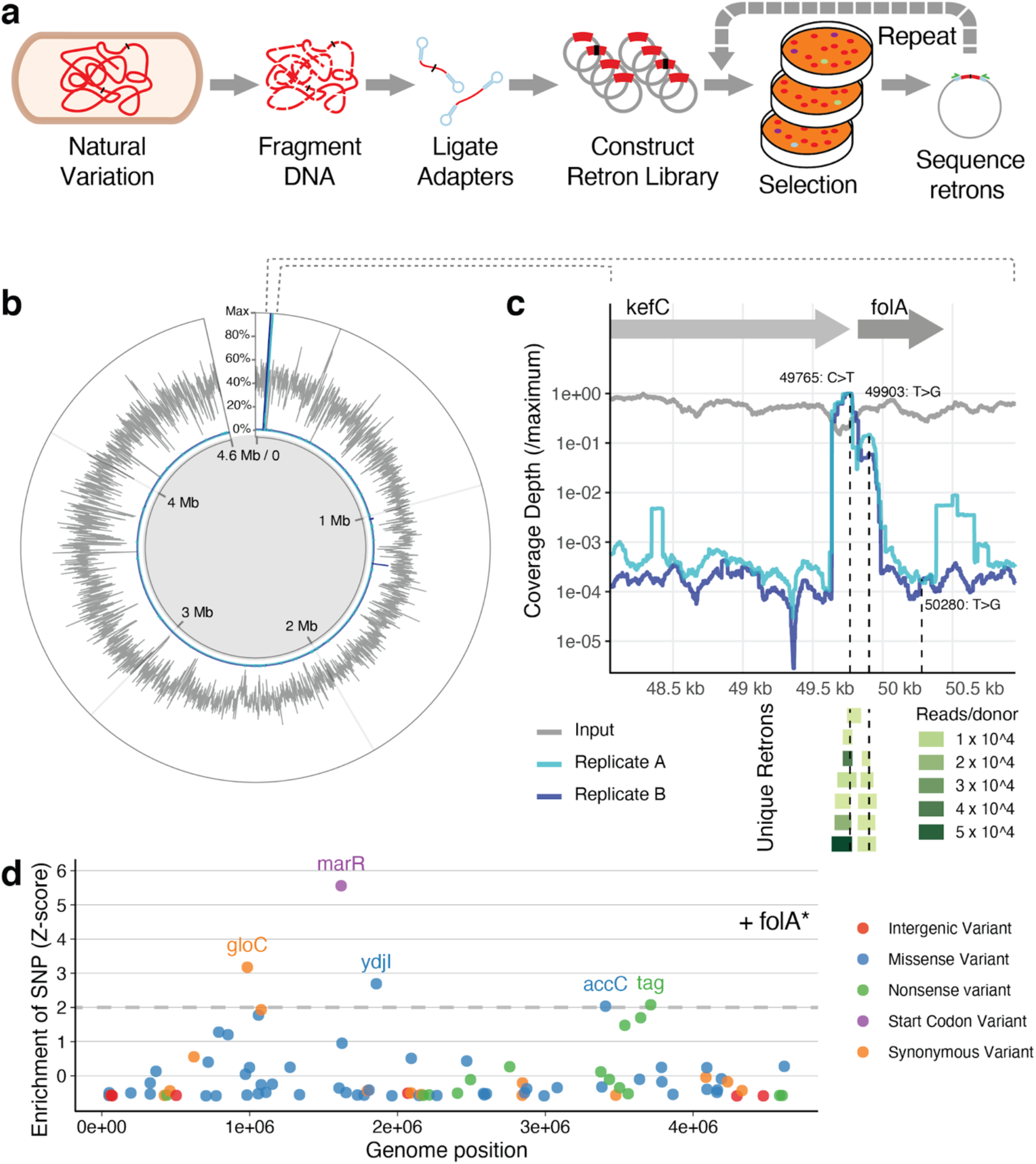
Detecting causal variants in genomic DNA using retron libraries. **5A** Graphical summary of an RLR experiment with genomic DNA (gDNA) as a source of alleles. gDNA of an evolved strain is randomly fragmented, and identical adapters ligated to the ends of these fragments. These adapters allow pooled cloning, creating a library of retron plasmids having millions of members. This library is induced, a selection is conducted, and retron donors of surviving mutants are sequenced as a pool via amplicon NGS. Optionally, mutants surviving selection can be transformed by the pool again, screening for additional mutations and combinatorial effects. **5B** Results of genomic DNA screen. Coverage of unique donor sequences across all genomic positions is displayed for a Genomic Retron Library, showing the mean coverage across 1000 base-pair windows and normalizing to the maximum coverage observed (Gray, see Supplementary figure 6B for additional detail). After selection, the depth at which variants are observed in surviving retron donors is depicted for two replicates of retron induction and selection on Trimethoprim, normalized to the maximum depth observed at a genomic position(blue, light blue). **5C** The region surrounding the *folA* locus is displayed, with genomic position on the X axis. Sequence coverage observed for each base is plotted on the Y axis for the input genomic retron library and for two replicates post-selection, relative to the maximum coverage observed in this region for a given sample. Below, retron donor sequences observed more than 1000 times in replicate A are depicted as a “pileup” aligned to the genome, and are colored by their abundance in post-selection sequencing. SNPs detected at the *folA* locus are indicated by vertical dashed lines, highlighting enrichment of two SNPs, and lack of enrichment of a third. **5D** Transforming the library into a strain already bearing these detected *folA* mutations (*folA\**) and plating on additional Trimethoprim exposes additional candidate causal variants. Retron donors sequenced post-selection are used to calculate Z-scores for each allele, describing their deviation from mean allele depth of coverage. Variants with Z-scores over 2 have been labeled by the gene in which they occur, and variants have been classified by their relationship to coding sequences.

Induction of RLR and selection with Trimethoprim dramatically increased the abundance of some donors containing variants at the folA locus (Fig 4B, 4C), which encodes the protein target of TMP^30^. Within RLR donors mapping to this region, coverage of two SNPs are highly enriched, indicating they individually increase TMP resistance (Fig. 4C). Multiple retron donor sequences independently contribute to the enrichment of both alleles (Fig. 4C). The more highly-enriched allele lies upstream of the folA CDS and likely increases it’s transcription, a well-described route to TMP resistance^31^. The other is known to increase the catalytic rate of FolA, leading to TMP resistance^31^. A third SNP within folA is not enriched, suggesting little or no effect individually at this concentration of TMP. Mutations at this third position are known to interact with folA mutations at the active site, but not confer resistance on their own^32^. The ability to individually measure mutational effect during the selection itself, especially for mutations so near each other in the genome, distinguishes this method from existing techniques like P1 transduction, which transfer up to ~100kb of contiguous variation, and require subsequent sequencing of hits for interpretation^33^.

To identify alleles outside the folA locus providing additional resistance, we performed a subsequent transformation of the genomic library into a strain bearing all folA mutations. Sequencing of retron plasmids following selection with increased TMP identified several enriched variants (Fig. 4D), including variants within known resistance determinants such as the marR regulator of the multi-antibiotic resistance operon^34^. In this manner, causal alleles can be determined repeatedly from evolved or environmentally derived pools of variation, in order to deconvolute phenotypes requiring multiple mutations.

## Discussion

Here we show that retron recombineering is a flexible, generalizable tool for genome editing, surpassing the editing rates achieved by other markerless editing tools like oligonucleotide recombineering. Pooled, barcoded mutant libraries can be prepared in this way and used for multiplexed characterization of natural and synthesized allelic variants, a process we call RLR. RLR enables millions of experiments to be performed simultaneously, and genome-wide insights to be gained.

RLR is an alternative to CRISPR-based methods which also perform pooled phenotypic measurements of mutant sequences. These CRISPR-based methods create edits by using a guide RNA to direct targeted breakage, and a plasmid-borne donor DNA to repair these breaks with the desired sequence, with phenotypic tracking permitted by amplicon sequencing of these components^35–38^. RLR’s “donor only” nature builds upon these “Guide + Donor” methods in several key ways. RLR eliminates the requirement for a minimum edited difference from the reference sequence, including single-base pair changes, to be characterized without requiring additional synonymous mutations being incorporated solely to ablate recognition by a guide^35–37^. RLR also overcomes the requirement for targeting a suitable protospacer-adjacent motif (PAM), which reduces editing efficiency of CRISPR-based methods as the distance to a PAM increases^35–37^. Intriguingly, production of ssDNA using a retron appears to improve break-dependent methods in S. cerevisiae^35^ and E. coli^39^.

That RLR does not require a “guide” element paired with an editing donor simplifies RLR elements. In contrast to the two unique elements required for “guide + donor” strategies, or the three required for efficient Prime editing^40^, RLR’s sole requirement is one unique donor sequence within the retron. This relaxed design constraint enables RLR using non-designed variation. We demonstrate this using random fragments of evolved DNA as an input. We expect that this approach will also enable use of random, degenerate variation introduced into templates, which is of great interest for characterizing sequences in-depth, and creating synthetic evolution systems^41^.

That CRISPR is not a required component of RLR also creates the possibility that RLR could be combined with engineered selections/screens of which CRISPR is already a component. In addition, Cas9 is toxic in many bacteria, even when nuclease-null forms are used^42, 43^. Non-toxic editing methods may enable new applications which may otherwise be hampered by expression of deleterious components.

The continuous, replication-dependent nature of RLR over multiple generations makes it distinct from break-dependent methods which occur as a discrete process. This feature facilitates RLR’s high efficiency, but may limit its utility in organisms where it is not practical or desirable to undergo multiple generations during editing. Editing over multiple generations may confer unique benefits however, such as the ability to extend editing periods for higher efficiency editing, and ability to edit during selection regimes for directed evolution. Continuous editing is best understood through a population genetics framework, and we offer a straightforward model for simulating these processes.

We have shown that editing is functional and high-efficiency across a sample of donor sequences, but more work remains to understand variation in editing, and further improve editing rates. RLR could feasibly be applied in a combinatorial manner, characterizing groups of mutations, and strategies remain to be developed in this area. It also remains unexplored whether RLR can be applied to large deletions and insertions, and possibly facilitate other alterations such as inversions, duplications, and rearrangements.

Here RLR measures beneficial growth and selective phenotypes, and we outline how improvement of editing makes deleterious phenotypes accessible. Non-growth phenotypes could also be made accessible to pooled measurement by fluorescent reporters^44^, biosensors^45^, single-cell transcriptomics^46^, and a myriad of other screens^8^. Likewise, given that recombineering is possible in a range of organisms including Eukaryotes^46,47^, and retrons occur across a range of organisms^16^, RLR should not be limited to use in *E. coli*, and development of RLR in other genetic systems is an exciting area of future research.

## Supporting information

Supplemental Sequences, Tables

## Online Methods

### Preparation of Strains and plasmids

Strain ECNR1 was modified to replace the bla ampicillin resistance cassette and the tetR repressor at the lambda prophage locus with the tetA tetracycline resistance cassette, using double-stranded recombineering as per Datsenko and Wanner^1^. All strains were also modified by inactivation of the araBAD operon using this method, conferring arabinose auxotrophy and ensuring consistent induction with arabinose. Other genes were likewise inactivated where relevant, and the antibiotic resistance markers removed using FLP recombinase as previously described^2^. Retron plasmid pFF745 (Addgene #61450) was a generous gift from Fahim Farzadfard and Timothy Lu. New retron donor sequences were introduced by KLD mutagenesis (New England Biolabs, NEB), or Gibson Assembly (NEB). Plasmids were modified to contain a low-copy origin under stringent replication control (SC101) and a tightly-regulated promoter (AraC-pBAD) to minimize variability in growth and plasmid titer among retron plasmid-bearing populations(Fig S5), and enable tight repression of the editing system. Expression in this system produced an effect on growth comparable to expressing GFP, an improvement on pFF745-derived COLE1-pL_lacO expression systems which had larger impacts on growth(Fig S5). Beta recombinase was replaced by CspRecT and mutL-E32K was added in relevant plasmids by Gibson Assembly (NEB). See Supplemental Sequences for oligonucleotides used to perform these alterations.

### Measurement of Editing

To perform editing, strains were transformed with retron plasmids via electroporation^1^, and plated to Lysogeny Broth (LB, Lennox formulation: 10g tryptone, 5g yeast extract, 5g sodium chloride per 1 liter distilled, deionized water, or ddH2O) with 25ug/mL Chloramphenicol(LB-CM25) with 1.5% agar added. After 18-24 hours of growth at 34°C, colonies were picked into 100uL of LB-CM25 in a 96-well plate and allowed to grow 6-8 hours, reaching confluence. These un-induced pre-cultures were diluted 1000-fold into 1mL of LB-CM25 containing 0.2% L-arabinose (LB-CM25ara) in a 96-well plate, and allowed to grow for 24 hours at 34°C with shaking at 900rpm, reaching confluence. This 1000-fold dilution and growth was repeated once more for all cultures. Assuming density at confluence to be consistent, the number of generations experienced with induction across both 1:1000 batch cultures is approximately 20, because 2^10 is equal to 1024. 120uL of these saturated cultures (OD_600_ approximately 3.0) were sampled to measure editing via amplicon sequencing as described below.

When growth in a tubidostat is indicated, strains were grown in an eVOLVER instrument^2^ (FynchBio). Turbidostat vials were maintained between OD_600_ of 0.2 and 0.4, in LB-CM25ara with 0.05% tween added to prevent biofilm formation, growing at 34 degrees with a stir rate of 8. 1mL samples were sampled at the indicated time points to measure editing via amplicon sequencing as described below.

To measure editing after batch growth, samples were centrifuged for 5 minutes at 4.8krcf, the pellet resuspended in 10mM NaOH solution with 0.01% Triton-100, and incubated at 95°C for 10 minutes in a thermocycler. The resulting lysates were centrifuged for 10 minutes at 4.8krcf at 4 degrees, and supernatants were stored at −20°C. PCR amplification of the genomic region was performed in reactions containing 10uL Q5 2× Mastermix (NEB), 2uL lysate supernatant, 0.2uL 50uM primer mixture for the locus (Supplemental sequences), 1uL Evagreen dye (Biotium), and 6.8uL ddH2O. Amplification was monitored in the SYBR channel on an Eppendorf Realplex4 real-time PCR system until several cycles of productive amplification were observed (typically less than 10 cycles), and the resulting amplicons were indexed for sequencing as described below.

### Preparation of genome-derived retron plasmid libraries

Trimethoprim-adapted strains s100 and s102 were a kind gift from Michael Baym^4^, MG1655 was obtained from the Coli Genetic Stock Center (CGSC, Yale University), and genomic DNA was isolated from 1.5mL LB cultures using the Promega Wizard Genomic DNA purification kit (Promega), and quantified using the Qubit dsDNA HS reagent (Thermo Fisher). 4ug DNA was placed in a Covaris 520045 tube, and sheared for 1200 seconds in a Covaris M220 using 50W peak incident power, 20% duty factor, and 200 cycles per burst. Fragmented DNA was visualized with a Bioanalyzer (Agilent), using a DNA 1000 chip (Agilent).

The resulting fragments were repaired, dA-tailed, and ligated to a custom adapter sequence (see Supplemental sequences) using the NEBnext Ultra II DNA library prep kit for Illumina (NEB). The resulting purified products were amplified in a reaction containing 25uL Q5 2× Mastermix (NEB), 2uL template, 0.5uL 50uM primer mixture (Supplemental sequences), 2.5uL Evagreen dye (Biotium), and 22.5uL ddH2O. Amplification was monitored in the SYBR channel on an Eppendorf Realplex4 real-time PCR system until several cycles of productive amplification were observed, typically less than 10 cycles, and were purified using DNA-binding magnetic beads^3^.

Adapter-dimer fragments containing no ligated insert were removed by digestion with BsaI enzyme and size-selective purification using DNA-binding magnetic beads^3^. After this purification, adapter-dimer fragments were no longer detectable by gel-electrophoresis, indicating their rarification in the pooled library.

A vector for Type-IIs “Golden Gate” assembly was prepared via PCR, using a pBAD-SC101 retron plasmid (with Beta or CspRecT, as indicated) as a template, and the resulting amplicon was purified using DNA-binding magnetic beads^3^. Vector and insert were ligated together in a Type-IIs “Golden Gate” assembly, with restriction and ligation occurring in one reaction mixture. This was performed in a reaction containing approximately .3pMol vector and 1pMol inserts, 2uL BsaI-v2, 2uL concentrated T4 DNA ligase (M202T, NEB). Reactions were incubated for 5 minutes at 37degC, then for 30 cycles of 37degC and 20degC for 4 minutes each, then finally for 60degC for 10 minutes, in a thermocycler.

### Preparation of Synthetic retron plasmid libraries

Recombineering donors were designed using the MODEST web tool^4^. Donors were assembled from individual oligonucleotides (Integrated DNA Technologies) using Primerize^5^. These donors were assembled into a compatible vector prepared by PCR (see supplemental sequences).

### Introduction of plasmid libraries into strains, editing

Plasmid library assembly reactions were purified by Ethanol precipitation, eluted into 2uL of TE buffer, and chilled on ice. 50uL of thawed electrocompetent cells (Lucigen ELITE 10G) were introduced, and electroporated using the EC1 setting on a Bio-Rad MicroPulser. After recovery in Lucigen Recovery medium for 1 hour at 37degC with shaking, 10mL of LB with 25ug/mL chloramphenicol was added for overnight growth at 30degC with shaking. Glycerol stocks were prepared at this point, and plasmids were isolated by midi-prep(Qiagen).

Electrocompetent cells were prepared by growing 50mL of Δ*mutS*, Δ*recJ*, Δ*sbcB* in LB at 30degC until an OD_600_ of 0.5 to 0.8 was achieved. These cultures were chilled on ice; pelleted, resuspended, and rinsed twice with 50mL chilled 10% glycerol; then resuspended in 2mL chilled glycerol for a final centrifugation. This pellet was resuspended in 150uL 10% glycerol, and 50uL aliquots were used for electroporation and recovery as described above.

This recovery was washed with PBS, resuspended into LB and a 1/5 dilution of this recovery was performed into 10mL LB-CM25Ara and grown for 6-8 hours at 34degC to saturate the culture and dilute out non-transformed and/or dead cells from the recovery. Editing was performed by diluting 1/1000 into 10mL(Synthetic libraries) or 50mL (genomic libraries) of LB-CM25Ara and grown to saturation 34degC overnight. This process was repeated once more to achieve greater than 20 generations of growth on average with induction.

### Binary Screens for drug resistance

Barcoded Synthetic mutant libraries prepared as above were pelleted and rinsed with PBS, before resuspending in PBS. Diluted samples were plated onto 100mm petri dishes containing LB with relevant concentrations of rifampicin and 1ug/mL chloramphenicol, to determine the CFU/ml plating efficiency across different conditions when grown overnight at 34degC. Not fewer than 6 petri dishes per condition were plated using a dilution targeting one thousand colonies per plate, and grown overnight at 34degC. The resulting colonies were scraped from the plates, rinsed and resuspended in PBS, and plasmids were isolated by Mini-prep (Qiagen).

For genomic libraries, the identical procedure was performed, except 150mm petri plates containing Cation-adjusted Miller-Hinton Broth containing 1ug/mL Trimethoprim and 1ug/mL Chloramphenicol were used, plates were incubated for 2 days at 34degC, and plasmids were isolated by Midi-prep (Qiagen). For the secondary screen in the folA* strain background, Trimethoprim was used at 1mg/mL, necessitating 1% DMSO in the medium for solubility.

### Quantitative Screen for drug resistance

Barcoded Synthetic mutant libraries prepared as above were diluted 1/1000 into 50mL of LB medium containing 10ug/mL Chloramphenicol and 5ug/mL rifampicin. After growth overnight at 34degC, a 5mL samples was taken, and this dilution was repeated for a second time point. Samples of the initial plasmid library, the library when transformed into Δ*mutS*, Δ*recJ*, Δ*sbcB* cells, and both timepoints, were obtained for plasmid purification.

### Deep Sequencing of Retron Donors

*E. coli* containing retron plasmids were obtained from the relevant liquid or solid growth condition, washed once with Dulbecco’s Phosphate-buffered Saline, and their plasmids purified via Miniprep or Midiprep (Qiagen). PCR amplification of retron donor sequences was performed in a reaction containing 20uL q5 2× Mastermix (NEB), 2uL purified plasmid DNA, 0.4uL 50uM primer mixture (Supplemental sequences), 1uL Evagreen dye (Biotium), and 15.6uL ddH2O. Amplification was monitored in the SYBR channel on an Eppendorf Realplex4 real-time PCR system until several cycles of productive amplification were observed, typically less than 10 cycles. Products were indexed for sequencing as described below.

### Amplicon sequencing

Amplicons from previous steps were prepared for Illumina sequencing by first removing oligonucleotides via treatment with Exonuclease I (NEB), then performing PCR using primers adding Indexes 1 and 2 for Illumina Paired-end sequencing(Supplemental sequences). PCR was performed in a reaction containing 10uL Q5 2X Mastermix (NEB), 1uL PCR product, .2uL 50uM primer mixture, 1uL Evagreen dye (Biotium), and 7.8uL ddH2O. Amplicons were purified using DNA-binding beads^3^ and quantified using the Qubit dsDNA HS reagent (Thermo Fisher). The resulting DNA were pooled and sequenced on MiSeq, NextSeq, or HiSeq Illumina instruments, producing paired-end, 150bp reads.

### Sequence Analysis: Editing

No fewer than 10,000 paired-end reads per replicate were merged using PEAR^6^ and adapter and primer sequences trimmed using Cutadapt^7^. Sequences surrounding the edited region were trimmed again with Cutadapt and counts of identical sequences were determined. Any sequence occurring fewer than 20 times was assumed to be a rare sequencing error, and was discarded at this stage. The edited fraction was calculated as the fraction of edited sequences divided by the number of edited and wild type reference sequences detected within a sample. To minimize the impact of sequencing error on conclusions, additional synonymous mutations were incorporated along with the desired missense *gyrA* and *rpoB* alleles for Figure 1B, where edited events are often rare. This more unambiguously links an edited sequence to an editing event, and not to mutation or errors during PCR/sequencing. A subset of these genotypes were additionally replicated by a phenotypic test- the efficiency of plating on antibiotic(Fig S1C). The fraction of all observed sequences which were either edited or wild type was monitored, and exceeded 93% for the results in Figure 2B and exceeded 99% for the additional alleles in Figure 2D. Scripts are available at https://github.com/churchlab/rlr.

### Culture-based confirmation of editing results

In parallel with amplicon sequencing to measure editing, serial dilutions of samples bearing the *RpoB* donor were plated to LB with 25ug/mL Chloramphenicol, and LB with 25ug Rifampicin. CFU/mL was determined by counting colonies, and the proportion of Rifampicin resistant CFU was used as a proxy for the edited fraction of cells.

### Sequence Analysis: Synthetic DNA experiments

Paired-end reads were merged using PEAR^6^, adapter and primer sequences were trimmed off using Cutadapt^7^, and counts of identical sequences were determined. Donor sequences not matching expected sequences were discarded. The fraction of sequences discarded correlated with estimated sequencing error rate and quality scores, and was never ≥25% of sequences. The frequency of each donor in the resulting dataset is then determined. For timecourses, these frequencies are normalized to the median frequency of the neutral allele retron pool within each replicate and time point, and initialized to their relative frequency in the first time point prior to plotting. For calculating relative growth rates, an exponential curve was fit to non-initialized trajectories using a linear model. Scripts are available at https://github.com/churchlab/rlr.

### Sequence Analysis: Natural DNA experiments

For determining the length of retron donor regions in cloned gDNA libraries, reads were first aligned to the MG1655 reference genome using BWA-MEM^8^. The length of the region between aligned read pairs, the Template Length or Tlen column, was extracted from this alignment and visualized as a histogram(Fig S6A). Template lengths longer than 500 base pairs were assumed to be the result of erroneous alignment, and were discarded.

For other analyses paired-end reads were merged using PEAR^6^ and adapter and primer sequences trimmed off using Cutadapt^7^. The resulting sequences were aligned to the MG1655 reference genome using BWA-MEM^8^. To determine coverage of the reference genome, depth was determined using Bedtools genomecov^9^, and visualized by plotting the mean coverage of 1000bp sliding windows(Fig 4B). Alternatively, to examine non-redundant coverage conferred by unique donors, sequences with identical start and end alignment positions were collapsed into a single sequence before determining depth, visualized by plotting the mean coverage of 500bp windows(Fig S6B). When examining a subset of the genome, coverage at every base was used, rather than using sliding windows (Fig 4C).

To determine the abundance of retron donors conferring SNPs, reads from amplicons obtained after selection were aligned to the MG1655 reference genome and SNPs inferred using Millstone^10^. Depth of SNP coverage in the resulting output reflects abundance in the pool, and was used for visualization and analysis.

### Individual Growth rate measurements

Colonies were picked into 100uL of LB medium in 96 well plates and grown for 4-6 hours at 34 degrees. 10uL of a 10-fold dilution of this “pre-culture” was used to inoculate 190uL of LB medium with relevant additives in a Nunclon 96-well microwell plate (Thermo Scientific). These plates were cultured in a shaking plate reader at 34 degrees, measuring OD_600_ every 15 minutes. The resulting data were analyzed using the analyze_growth.m script to derive growth rates for all wells. Scripts are available at https://github.com/churchlab/rlr.

### Population genetics model

A simulation was written in the Matlab scripting language, in which each generation, a fraction ‘r’ of un-edited cells become edited, for a population bearing a particular retron. Once edited, that population reproduces at a rate of ‘f’, which may be more or less than the parental rate of 1. Simulations can be performed with various r and f and initial conditions. The model makes implicit assumptions that populations experience constant, uniform exponential growth and never approach carrying capacity of their environment. Scripts are available at https://github.com/churchlab/rlr.

## References

1. Baba, T. et al. Construction of Escherichia coli K-12 in-frame, single-gene knockout mutants: the Keio collection. Mol. Syst. Biol. 2, 2006.0008 (2006).

2. Richardson, S. M. et al. Design of a synthetic yeast genome. Science 355, 1040–1044 (2017).

3. Hutchison, C. A., 3rd et al. Design and synthesis of a minimal bacterial genome. Science 351, aad6253 (2016).

4. van Opijnen, T. & Camilli, A. Transposon insertion sequencing: a new tool for systems-level analysis of microorganisms. Nat. Rev. Microbiol. 11, 435–442 (2013).

5. Warner, J. R., Reeder, P. J., Karimpour-Fard, A., Woodruff, L. B. A. & Gill, R. T. Rapid profiling of a microbial genome using mixtures of barcoded oligonucleotides. Nat. Biotechnol. 28, 856–862 (2010).

6. Gilbert, L. A. et al. Genome-Scale CRISPR-Mediated Control of Gene Repression and Activation. Cell 159, 647–661 (2014).

7. Peters, J. M. et al. A Comprehensive, CRISPR-based Functional Analysis of Essential Genes in Bacteria. Cell 165, 1493–1506 (2016).

8. Gray, A. N. et al. High-throughput bacterial functional genomics in the sequencing era. Curr. Opin. Microbiol. 27, 86–95 (2015).

9. Tenaillon, O. et al. Tempo and mode of genome evolution in a 50,000-generation experiment. doi:10.1101/036806.

10. Heim, R. & Tsien, R. Y. Engineering green fluorescent protein for improved brightness, longer wavelengths and fluorescence resonance energy transfer. Curr. Biol. 6, 178–182 (1996).

11. Hong, K.-K. & Nielsen, J. Recovery of phenotypes obtained by adaptive evolution through inverse metabolic engineering. Appl. Environ. Microbiol. 78, 7579–7586 (2012).

12. Costantino, N. & Court, D. L. Enhanced levels of lambda Red-mediated recombinants in mismatch repair mutants. Proc. Natl. Acad. Sci. U. S. A. 100, 15748–15753 (2003).

13. Nyerges, Á. et al. A highly precise and portable genome engineering method allows comparison of mutational effects across bacterial species. Proc. Natl. Acad. Sci. U. S. A. 113, 2502–2507 (2016).

14. Wang, H. H. et al. Programming cells by multiplex genome engineering and accelerated evolution. Nature 460, 894–898 (2009).

15. Lampson, B. C., Inouye, M. & Inouye, S. Retrons, msDNA, and the bacterial genome. Cytogenet. Genome Res. 110, 491–499 (2005).

16. Simon, A. J., Ellington, A. D. & Finkelstein, I. J. Retrons and their applications in genome engineering. Nucleic Acids Res. (2019) doi:10.1093/nar/gkz865.

17. Farzadfard, F. & Lu, T. K. Synthetic biology. Genomically encoded analog memory with precise in vivo DNA writing in living cell populations. Science 346, 1256272 (2014).

18. Lajoie, M. J., Gregg, C. J., Mosberg, J. A., Washington, G. C. & Church, G. M. Manipulating replisome dynamics to enhance lambda Red-mediated multiplex genome engineering. Nucleic Acids Res. 40, e170 (2012).

19. Mosberg, J. A., Lajoie, M. J. & Church, G. M. Lambda red recombineering in Escherichia coli occurs through a fully single-stranded intermediate. Genetics 186, 791–799 (2010).

20. Sawitzke, J. A. et al. Probing cellular processes with oligo-mediated recombination and using the knowledge gained to optimize recombineering. J. Mol. Biol. 407, 45–59 (2011).

21. Mosberg, J. A., Gregg, C. J., Lajoie, M. J., Wang, H. H. & Church, G. M. Improving Lambda Red Genome Engineering in Escherichia coli via Rational Removal of Endogenous Nucleases. PLoS ONE vol. 7 e44638 (2012).

22. Simon, A. J., Morrow, B. R. & Ellington, A. D. Retroelement-Based Genome Editing and Evolution. ACS Synth. Biol. 7, 2600–2611 (2018).

23. Farzadfard, F., Gharaei, N., Citorik, R. J. & Lu, T. K. Efficient Retroelement-Mediated DNA Writing in Bacteria. doi:10.1101/2020.02.21.958983.

24. Wannier, T. M. et al. Improved bacterial recombineering by parallelized protein discovery. doi:10.1101/2020.01.14.906594.

25. Lajoie, M. J. et al. Genomically recoded organisms expand biological functions. Science 342, 357–360 (2013).

26. Barrick, J. E., Kauth, M. R., Strelioff, C. C. & Lenski, R. E. Escherichia coli rpoB Mutants Have Increased Evolvability in Proportion to Their Fitness Defects. Molecular Biology and Evolution vol. 27 1338–1347 (2010).

27. Pozzi, G. et al. rpoB mutations in multidrug-resistant strains of Mycobacterium tuberculosis isolated in Italy. J. Clin. Microbiol. 37, 1197–1199 (1999).

28. Tenaillon, O. et al. The Molecular Diversity of Adaptive Convergence. Science vol. 335 457–461 (2012).

29. Kepler, T. B. & Perelson, A. S. Drug concentration heterogeneity facilitates the evolution of drug resistance. Proc. Natl. Acad. Sci. U. S. A. 95, 11514–11519 (1998).

30. Baym, M. et al. Spatiotemporal microbial evolution on antibiotic landscapes. Science 353, 1147–1151 (2016).

31. Tamer, Y. T. et al. High-Order Epistasis in Catalytic Power of Dihydrofolate Reductase Gives Rise to a Rugged Fitness Landscape in the Presence of Trimethoprim Selection. Mol. Biol. Evol. 36, 1533–1550 (2019).

32. Dion, A., Linn, C. E., Bradrick, T. D., Georghiou, S. & Howell, E. E. How do mutations at phenylalanine-153 and isoleucine-155 partially suppress the effects of the aspartate-27-->serine mutation in Escherichia coli dihydrofolate reductase? Biochemistry 32, 3479–3487 (1993).

33. Thomason, L. C., Costantino, N. & Court, D. L. E. coli genome manipulation by P1 transduction. Curr. Protoc. Mol. Biol. Chapter 1, Unit 1.17 (2007).

34. Sulavik, M. C., Gambino, L. F. & Miller, P. F. The MarR repressor of the multiple antibiotic resistance (mar) operon in Escherichia coli: prototypic member of a family of bacterial regulatory proteins involved in sensing phenolic compounds. Molecular Medicine 1, 436 (1995).

35. Sharon, E. et al. Functional Genetic Variants Revealed by Massively Parallel Precise Genome Editing. Cell 175, 544–557.e16 (2018).

36. Garst, A. D. et al. Genome-wide mapping of mutations at single-nucleotide resolution for protein, metabolic and genome engineering. Nat. Biotechnol. 35, 48–55 (2017).

37. Roy, K. R. et al. Multiplexed precision genome editing with trackable genomic barcodes in yeast. Nat. Biotechnol. 36, 512–520 (2018).

38. Guo, X. et al. High-throughput creation and functional profiling of DNA sequence variant libraries using CRISPR–Cas9 in yeast. Nature Biotechnology vol. 36 540–546 (2018).

39. Lim, H. et al. Multiplex generation, tracking, and functional screening of substitution mutants using a CRISPR/retron system. doi:10.1101/2020.01.02.892679.

40. Anzalone, A. V. et al. Search-and-replace genome editing without double-strand breaks or donor DNA. Nature vol. 576 149–157 (2019).

41. Simon, A. J., d’Oelsnitz, S. & Ellington, A. D. Synthetic evolution. Nat. Biotechnol. 37, 730–743 (2019).

42. Cho, S. et al. High-Level dCas9 Expression Induces Abnormal Cell Morphology in Escherichia coli. ACS Synth. Biol. 7, 1085–1094 (2018).

43. Rock, J. M. et al. Programmable transcriptional repression in mycobacteria using an orthogonal CRISPR interference platform. Nat Microbiol 2, 16274 (2017).

44. Kosuri, S. et al. Composability of regulatory sequences controlling transcription and translation in Escherichia coli. Proc. Natl. Acad. Sci. U. S. A. 110, 14024–14029 (2013).

45. Rogers, J. K. et al. Synthetic biosensors for precise gene control and real-time monitoring of metabolites. Nucleic Acids Res. 43, 7648–7660 (2015).

46. Dixit, A. et al. Perturb-Seq: Dissecting Molecular Circuits with Scalable Single-Cell RNA Profiling of Pooled Genetic Screens. Cell 167, 1853–1866.e17 (2016).

47. Murphy, K. C. λ Recombination and Recombineering. EcoSal Plus vol. 7 (2016).

48. Barbieri, E. M., Muir, P., Akhuetie-Oni, B. O., Yellman, C. M. & Isaacs, F. J. Precise Editing at DNA Replication Forks Enables Multiplex Genome Engineering in Eukaryotes. Cell 171, 1453–1467.e13 (2017).

## Online Methods References

1. Datsenko, K. A. & Wanner, B. L. One-step inactivation of chromosomal genes in Escherichia coli K-12 using PCR products. Proc. Natl. Acad. Sci. U. S. A. 97, 6640–6645 (2000).

2. Wong, B. G., Mancuso, C. P., Kiriakov, S., Bashor, C. J. & Khalil, A. S. Precise, automated control of conditions for high-throughput growth of yeast and bacteria with eVOLVER. Nat. Biotechnol. 36, 614–623 (2018).

3. Oberacker, P. et al. Bio-On-Magnetic-Beads (BOMB): Open platform for high-throughput nucleic acid extraction and manipulation. PLOS Biology vol. 17 e3000107 (2019).

4. Bonde, M. T. et al. MODEST: a web-based design tool for oligonucleotide-mediated genome engineering and recombineering. Nucleic Acids Res. 42, W408–15 (2014).

5. Tian, S. & Das, R. Primerize-2D: automated primer design for RNA multidimensional chemical mapping. Bioinformatics 33, 1405–1406 (2017).

6. Zhang, J., Kobert, K., Flouri, T. & Stamatakis, A. PEAR: a fast and accurate Illumina Paired-End reAd mergeR. Bioinformatics 30, 614–620 (2014).

7. Martin, M. Cutadapt removes adapter sequences from high-throughput sequencing reads. EMBnet.journal vol. 17 10 (2011).

8. Li, H. Aligning sequence reads, clone sequences and assembly contigs with BWA-MEM. arxiv.org (2013).

9. Quinlan, A. R. & Hall, I. M. BEDTools: a flexible suite of utilities for comparing genomic features. Bioinformatics 26, 841–842 (2010).

10. Goodman, D. B. et al. Millstone: software for multiplex microbial genome analysis and engineering. Genome Biol. 18, 101 (2017).

