## Supplemental Figures for "High throughput functional variant screens via in-vivo production of single-stranded DNA"

#### Supplemental Figures and materials

#### Supplemental Figures:

### Supplemental Figure 1

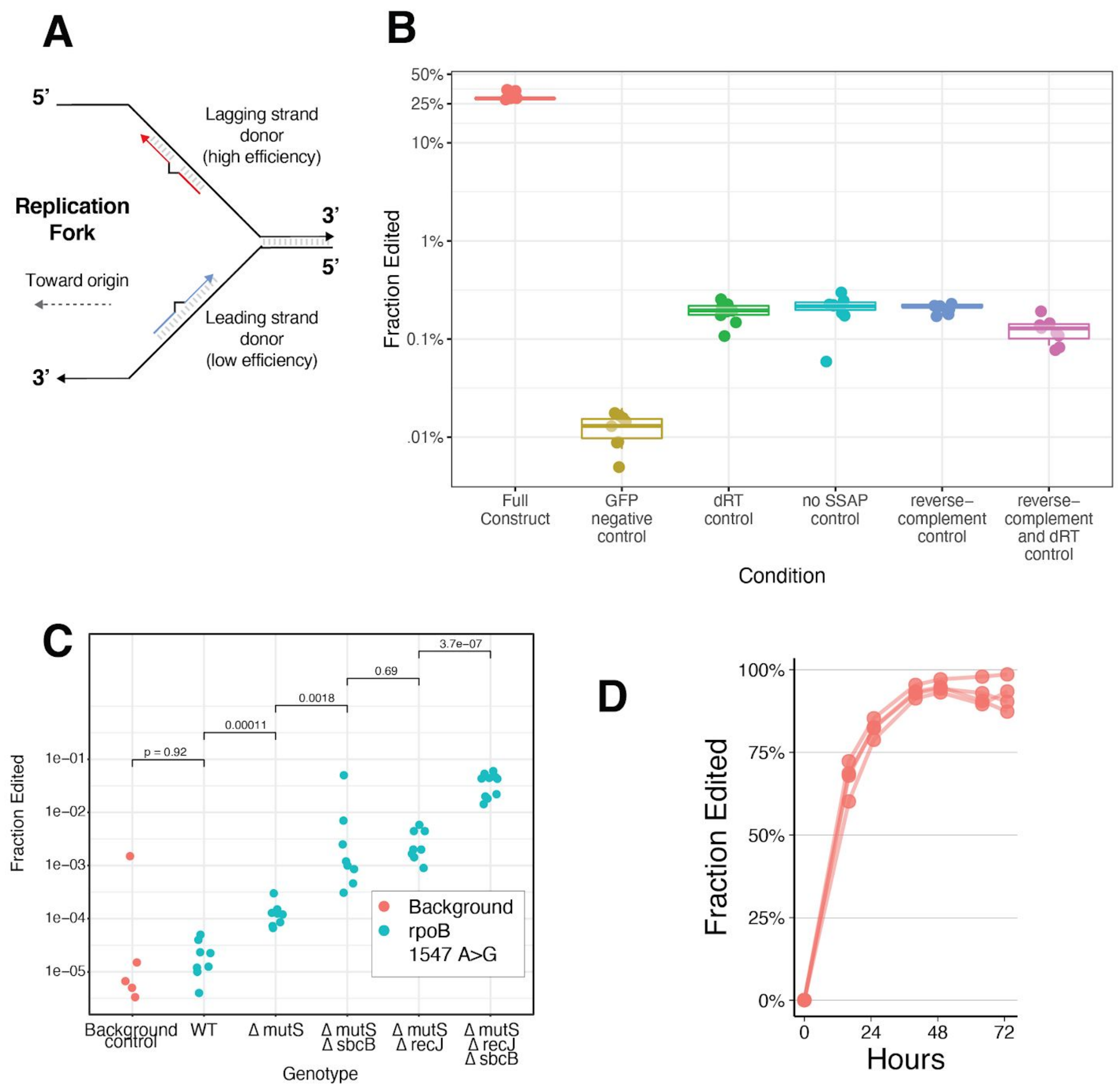

**Supplemental Figure 1:**

**S1A** A diagram of the replication fork, depicting recombineering donors annealing to either the leading strand of replication, or the lagging strand of replication. Recombineering donors annealing to the lagging strand typically edit more effectively[1].

**S1B** A confirmation of the mechanism of editing. Editing was compared between the full RLR system expressing the gyrA 248 C>T donor using Beta as a Single-stranded annealing protein (SSAP), and control constructs expressing only GFP, having an inactivated retron reverse-transcriptase active site (dRT[2]) and/or reverse-complemented donor DNA annealing to the leading strand, or which do not express an SSAP protein. Editing was performed in the ∆mutS ∆recJ ∆sbcB background, with Beta as an SSAP. Individual measurements are represented with dots, and the data is summarized by box plots in the style of Tukey.

**S1C** The edited fraction of cells, measured phenotypically by resistance to rifampicin at 25ug/mL. These results confirm those shown in Figure 1B, but are measured by plating for CFU, rather than by amplicon sequencing at the edited locus.

**S1D** The continuous nature of Retron editing. ∆mutS ∆recJ ∆sbcB cells expressing the gyrA retron with Beta as an SSAP were sampled during continuous growth and induction in a turbidostat. The edited fraction of the *gyrA* locus was determined by amplicon deep sequencing over time. This data also depicted in figure 2C, but with inferred generations of growth on the X axis.

#### Supplemental Figure 2

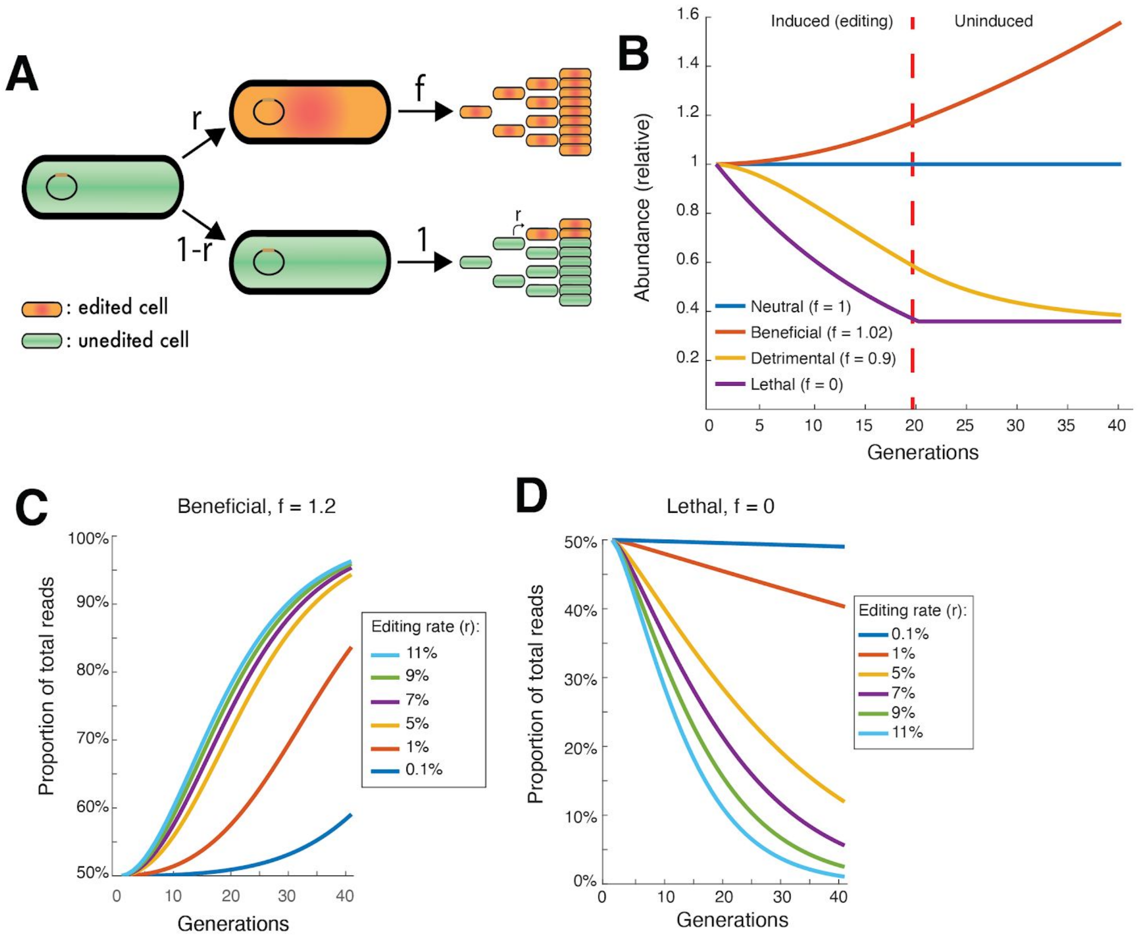


**Supplemental figure 2:**

**S2A** A population genetics model describing the Retron Library Recombineering (RLR) process. Each generation, a cell has probability "r" of being successfully edited by the retron construct it bears, and if so edited, propagates at rate "f" determined by the fitness of this mutation in future generations. If not edited in a given generation, which occurs at a rate 1-r, editing may occur in future generations.

**S2B** The population genetics model predicts that for beneficial alleles, the editing process can be halted (red vertical line), and the lineage of cells marked with the beneficial retron will continue to become more abundant. In contrast, for deleterious or lethal mutations, continued decrease in abundance of a barcoded population is dependent on continued induction and editing.

**S2C** The growth of populations bearing a retron conferring mutations with growth rate 120% that of the parent (f = 1.2) are simulated, for different rates of editing (r). Population size compared to a simulated neutral parent control, and thus proportion of barcode reads observed, is depicted on the Y axis. The signal observed for beneficial alleles is large, and does not vary substantially if editing frequency varies among alleles. This is compatible with quantitative measurement of beneficial allele fitness, even if editing rate varies somewhat. Additionally, beneficial alleles can be detected even with inefficient editing.

**S2D** Simulated growth of populations as in Figure S2C, except with a Lethal mutation (f = 0). In contrast with Figure S2C, the signal for deleterious or lethal alleles is comparatively small, and scales with editing rate to a greater degree. This indicates that quantification of deleterious phenotypes will be clouded by differences in editing rate among alleles to a greater degree, and that deleterious phenotypes require more efficient editing and/or editing over more generations for robust detection.

### Supplemental Figure 3

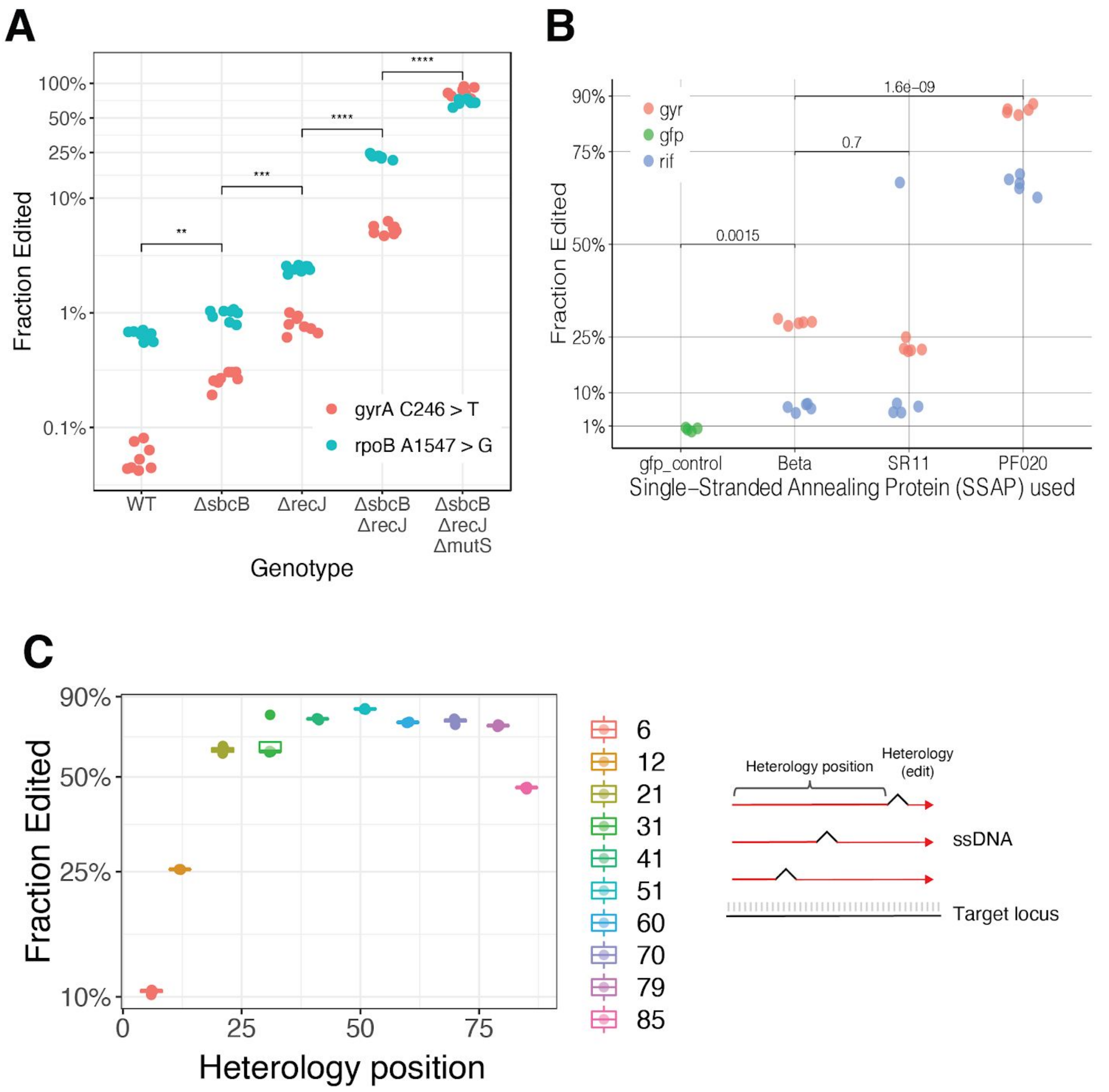


**Supplemental figure 3:**

**S3A** The effect of genotype on editing efficiency when the CspRecT is used rather than Beta as an SSAP. Individual measurements of the edited fraction of cells by amplicon sequencing are depicted by dots, for two loci. **** signifies $p < 0.0001$, *** signifies $p < 0.001$, and ** signifies $p < 0.01$, for two-tailed unpaired t-tests performed for comparisons indicated in brackets.

**S3B** The effect of SSAP protein on editing efficiency. Edited fraction of cells after batch growth was measured by amplicon sequencing for two loci. Retron constructs used Beta, "SR11", or CspRecT as their SSAP, or expressed only GFP as a negative control. Independent measurements are represented with dots, and p-values of two-tailed, unpaired t-tests are given for the comparisons shown.

**S3C** The effect of heterology position within a 90-mer donor DNA was determined for the gyrA 246 C>T mutation, using CspRecT as an SSAP. Donors conferring this mutation were constructed with the heterologous mutation present in different locations within the donor, as indicated in the diagram. Individual cultures were grown and induced for approximately 20 generations. Box plots in the style of Tukey are shown, and the 4 replicate measurements are additionally represented with dots.

### Supplemental Figure 4

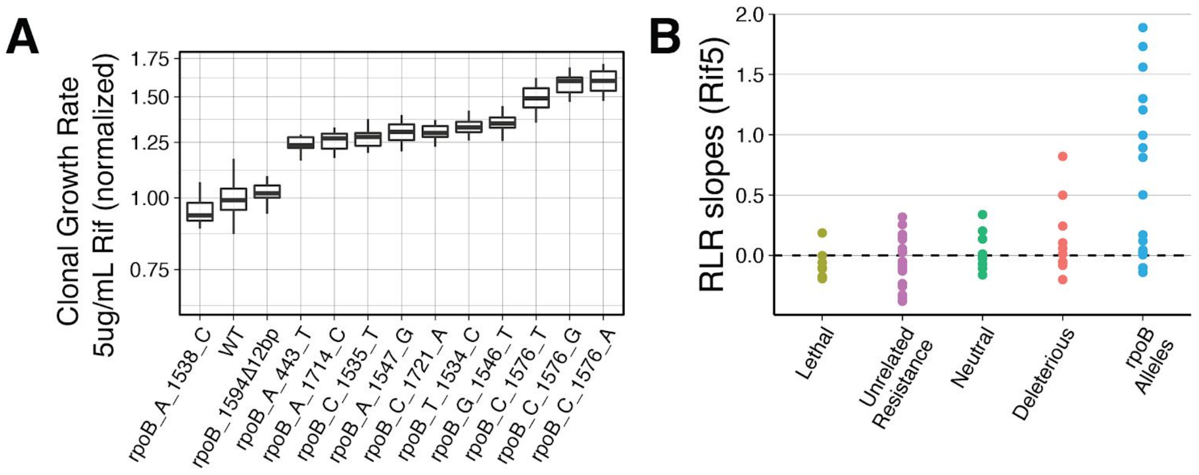


**Supplemental Figure 4**

**S4A** *rpoB* mutants conferring rifampicin resistance were constructed in the Δ*mutS* Δ*recJ* Δ*sbcB* background via oligonucleotide recombineering[3], and growth rates were measured in batch growth with sub-inhibitory rifampicin (5ug/ml). Data is shown normalized to the parental (WT, Δ*mutS* Δ*recJ* Δ*sbcB*) growth rate.

**S4B** Summary of RLR enrichment observed in a pooled, quantitative experiment over time. RLR enrichment rate (slope) for all alleles across 3 replicate experiments is indicated by a dot, and different classes of mutants are separated by color.

### Supplemental Figure 5

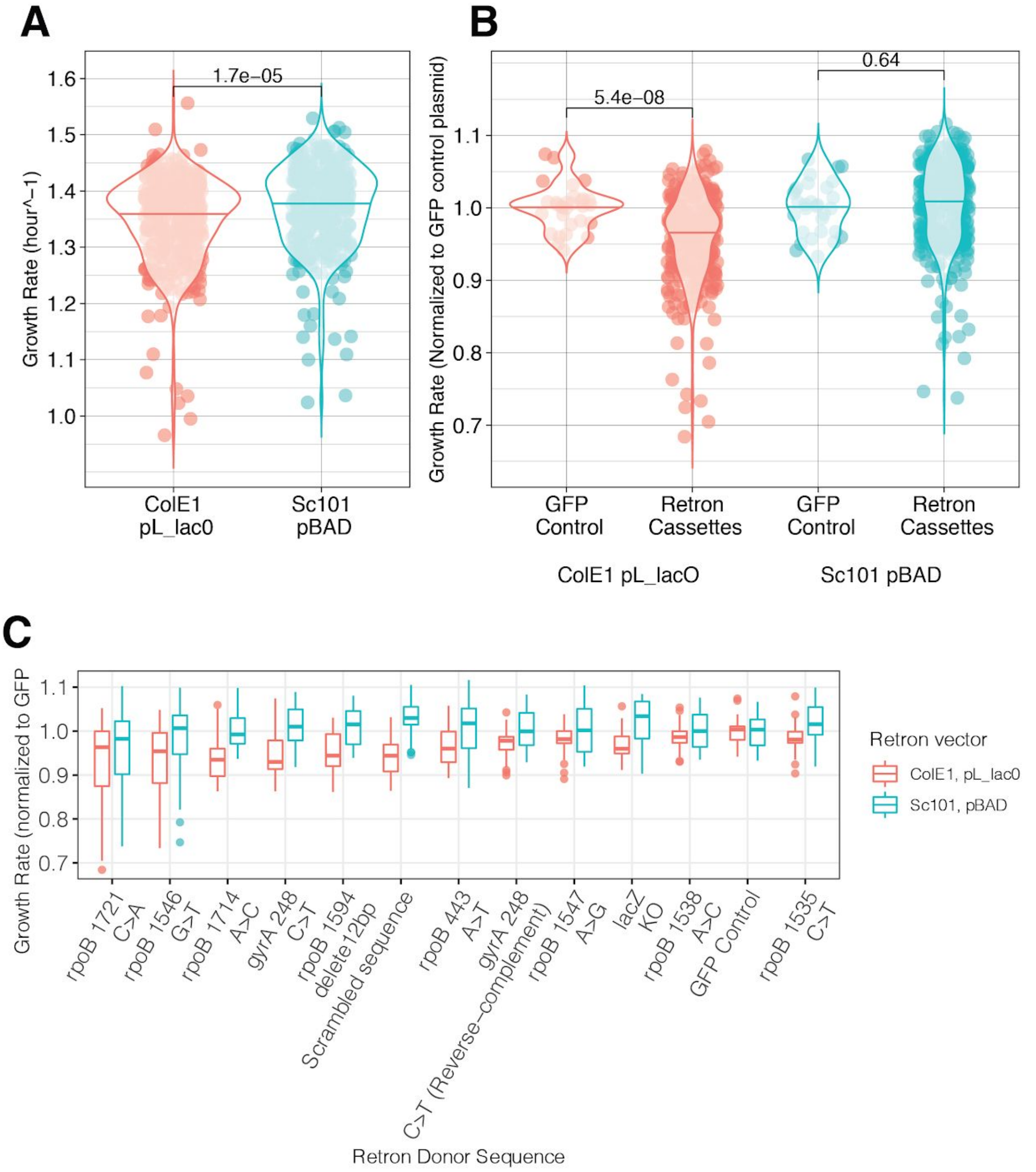


**Supplemental Figure 5**

**S5A** Where previous studies used a medium-copy ColE1 vector and pL_lacO promoter for expression of the retron components[1,2,4], expression from a low-copy (sc101) vector is less detrimental to growth. Absolute growth rates are pooled here for 12 expressed retron constructs, and violin plot outlines depict the distribution of the data, with the width of outlines indicating the proportion of the data with a given value. Horizontal lines depict the mean growth rate for each vector type. P value for a t-test comparing data for both vector types is given.

**S5B** Retrons are expressed from the ColE1 vector, they are significantly depressed in growth rate compared to sfGFP expressed in the same manner, whereas 12 retrons expressed from the SC101 vector cannot be distinguished, as a population. Growth rate is shown normalized to the GFP control construct for each vector type, and horizontal lines depict the mean growth rate for each condition. P values for t-tests are given for the comparisons indicated with brackets.

**S5C** Growth rates for 12 induced retron constructs, normalized to the GFP-expressing control plasmid for each vector type. Constructs are ordered by mean growth rate, and box-and-whisker plots are depicted in the style of Tukey, with outlier observations represented by dots.

### Supplemental Figure 6

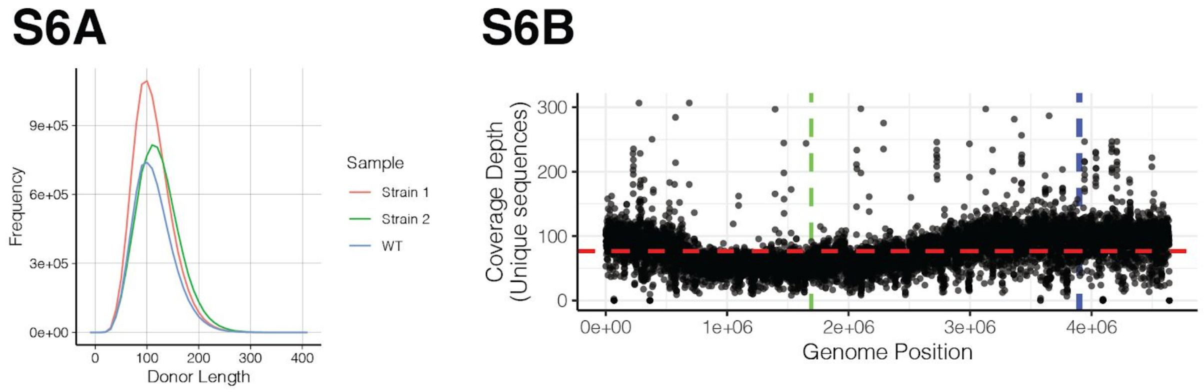


**Supplemental Figure 6**

**S6A** Retron donor size distribution for three representative genomic DNA libraries. Strains 1 and 2 were evolved, Trimethoprim-resistant *E. coli*, and "WT" is *E. coli* MG1655.

**S6B** Depth of coverage of a representative genomic retron library is depicted. Dots indicate the mean coverage of unique retron donor sequences across 500bp windows of the genome. The mean coverage across all windows is 78, and is indicated with a dashed red line. Dashed gray lines mark the genomic positions of the origin and terminus of the genome, helping to clarify increased coverage as DNA content increases with proximity to the origin. See Supplemental tables 4 and 5 for comments on regions of low and high coverage, respectively.

Supplemental Figures References:

1. Mosberg, J. A., Lajoie, M. J. & Church, G. M. Lambda red recombineering in Escherichia coli occurs through a fully single-stranded intermediate. *Genetics* **186**, 791–799 (2010).

2. Farzadfard, F. & Lu, T. K. Synthetic biology. Genomically encoded analog memory with

precise in vivo DNA writing in living cell populations. *Science* **346**, 1256272 (2014).

3. Wang, H. H. *et al.* Programming cells by multiplex genome engineering and accelerated evolution. *Nature* **460**, 894–898 (2009).

4. Simon, A. J., Morrow, B. R. & Ellington, A. D. Retroelement-Based Genome Editing and Evolution. *ACS Synth. Biol.* **7**, 2600–2611 (2018).

**Supplemental materials**:
All data and scripts necessary to reproduce figures and analysis can be found at https://github.com/churchlab/rlr

Index of supplemental materials:
**supplemental_sequences.xlsx**: annotated list of selected oligonucleotides used in the study.

**supplemental_sequence_maps**: generalized genbank plasmid maps for Retron Recombineering plasmids used in this study, and a sf.GFP-expressing control plasmid.

**Supplemental table 1**: summarized data from Figure 2B, showing the edited fraction measured by different genotypes
**Supplemental table 2**: summarized data from Figure 3C, showing the enrichment scores for all alleles

**Supplemental table 3**: summarized data from Figure 3D, showing enrichment scores for rpoB alleles across all rifampicin concentrations.

**Supplemental table 4**: regions of the RLR genomic Library for which Zero genomic coverage was observed. In all cases this was interpreted as artifacts due to the differences between the MG1655 reference genome and the BW25113::ΔlacA ancestor of the evolved strain. The region spanning the termini of the linear reference sequence also displayed artifactually low coverage when aligning sequences from the circular genome.

**Supplemental table 5**: regions of the RLR genomic Library for which very high coverage was observed. In all cases this was interpreted as artifacts due to mis-alignment of sequences present in multiple copies across the genome, such as insertion elements and ribosomal RNA.
